# Early viral infection of cyanobacteria drives bacterial chemotaxis in the oceans

**DOI:** 10.1101/2023.10.24.563588

**Authors:** Richard J. Henshaw, Jonathan Moon, Michael R. Stehnach, Benjamin P. Bowen, Suzanne M. Kosina, Trent R. Northen, Jeffrey S. Guasto, Sheri A. Floge

## Abstract

Interactions among marine microbes primarily occur through exudation and sensing of dissolved chemical compounds, which ultimately control ecosystem-scale processes such as biomass production, nutrient cycling, carbon fixation, and remineralization. Prior to lysis, viruses alter host metabolism, stimulating the release of dissolved chemical cues from intact plankton. However, the nature and degree of interactions between prelysis, virus-infected cells and neighbouring microbes remain unquantified. Here, we determine the impact of viral infection on dissolved metabolite pools from the marine cyanobacterium *Synechococcus* and the subsequent chemotactic response of heterotrophic bacteria using time-resolved metabolomics and microfluidics. Metabolites released from intact, virus-infected *Synechococcus* elicited vigorous chemoattractive responses from heterotrophic bacteria (*Vibrio alginolyticus* and *Pseudoalteromonas haloplanktis*), with the strongest responses occurring in the early infection stages and following cell lysis. We provide the first experimental observations of sustained chemotaxis towards live, infected *Synechococcus*, which is contrasted by no discernible chemotaxis toward uninfected *Synechococcus*. Finally, metabolite compounds and concentrations driving chemotactic responses were identified using a novel high-throughput microfluidic device. Our findings establish that prior to cell lysis, virus-infected picophytoplankton release compounds that significantly attract motile heterotrophic bacteria, illustrating a viable mechanism for resource transfer to chemotactic bacteria with implications for our understanding of carbon and nutrient flux across trophic levels.

## INTRODUCTION

Marine phytoplankton are responsible for half of global primary production [1], thus playing a critical role in ecosystem dynamics and biogeochemical cycles. Specifically, marine picocyanobacteria, including *Synechococcus* and *Prochlorococcus*, are the most abundant phototrophs on the planet. They contribute about one quarter of global oceanic primary production and dominate oligotrophic ocean surface waters, which are expected to increase in response to global warming [2]. Because of the diminutive stature of these micron-sized cells, their phycospheres - the chemical environment produced by and immediately surrounding phytoplankton cells - were long-thought to be too small to influence the surrounding chemotactic microbial community [3]. However, bacterial chemotaxis - directed motility in response to an external chemical gradient - towards extracellular compounds produced by *Synechococcus* and *Prochlorococcus* has previously been reported [4], and recent computational models [5] predict that chemotaxis may confer a substantial nutrient uptake advantage in cyanobacteria-bacteria interactions. Taken together, these works open the door to new mechanisms potentially promoting significant interactions across the complex network of marine microbes.

Viruses are abundant and dynamic components of marine microbial communities [6, 7], with marine cyanophage (viruses infecting cyanobacteria) regulating cyanobacterial mortality, community composition, and evolution [8]. Approximately ten percent of marine cyanobacteria are destroyed daily via viral lysis [6, 9–11], creating nutrient hotspots for chemotaxing bacteria that facilitate carbon and resource exchange [4, 12, 13]. While such lysis events are ephemeral and generally insufficient to support motile bacterial populations in non-bloom conditions [12], the viral infection cycle of host cells is prolonged over hours. Bacterial aggregation around intact, virus-infected microbes has been observed [14], during which viruses alter host cell metabolism [15] leading to both changes in metabolite composition and increased metabolite release from intact, infected cells [16– 18], including potential chemoattractants in the case of *Synechococcus* [19]. However, a dearth of direct experimental quantification persists. Our understanding of single-cell interactions between virus-infected cyanobacteria and surrounding microbes is limited, specifically how time-dependent changes in the phycosphere composition of infected cyanobacteria impact chemotaxis-mediated microbial interactions. Here, we quantify the time evolution of chemoattraction between virus infected *Synechococcus* and marine bacteria throughout an infection cycle. We interrogate a culture-based model system using time-resolved metabolomics to quantify virus-induced changes in cell exudates post infection compared to exudates derived from uninfected cells. Subsequently, microfluidics and video microscopy reveal the population-level response of two model chemotactic heterotrophic bacteria - *Vibrio alginolyticus* and *Pseudoalteromonas haloplanktis* - whose degree of chemoattraction to exudates from infected cyanobacteria evolves throughout the infection cycle versus non-infected cells. Using live phage infected *Synechococcus* cells, we further demonstrate that *V. alginolyticus* preferentially accumulates towards intact infected cells rather than uninfected host cells. Finally, a new high-throughput microfluidic platform is used to screen the chemotactic response of *V. alginolyticus* to specific exometabolites identified during various stages of viral infection. Our results establish how viral manipulation of host metabolism stimulates bacteria-cyanobacteria interactions with important implications for carbon and nutrient cycling.

## RESULTS

### Phage-induced changes in extracellular metabolites

Focusing on the model marine cyanobacteria *Synechococcus* WH8102, the impact of phage infection on extracellular nonpolar metabolite composition were determined over the course of an infection cycle with the lytic T4-like Myovirus S-SSM5. Phage infected and uninfected (control) cultures were centrifuged, washed, and resuspended in fresh medium after an initial phage adsorption period (four biological replicates each; see Methods). The onset of phage-induced cell lysis followed a *T* = 6 − 8 hpi (hours post infection) latent period. (Fig. 1a). Uninfected cultures exhibited no significant changes in cell density over the experimental time course. From the 980 metabolite features observed (857 in positive ionization mode, 123 in negative mode) during the infection cycle, a total of 7 exometabolites with functional annotations significantly changed (see Methods) in the phage-infected treatment as compared to the control uninfected treatment (Fig. 1bc, Extended Data Table I, Table S3) as determined by C18 chromatographic separations. Four compounds: 5’-methylthioadenosine, 5’-deoxyadenosine, L-phenylalanine, and palmitamide, were relatively enriched at most time points. Two compounds, dioctyl phthalate and tributylamine, exhibited depletion across all time points, with a third compound, N,N-dimethyldodeclamine N-oxide, depleted at both early (*T* = 2, 4.4 hpi) and late (*T* = 12 hpi) time points. Within the first 6 h post infection, four compounds were enriched in the extracellular fraction indicating pre-lysis metabolite release, and three compounds were depleted indicating either reduced release or increased uptake, relative to uninfected cells. Polar exometabolite analyses conducted on these same samples revealed 10 identified metabolites enriched in phage-infected cultures relative to the uninfected control and one metabolite depleted during the pre-lysis or latent phase of infection [19]. Thus, phage infection led to both increases and decreases of particular metabolites relative to uninfected cell cultures, and these exometabolite enrichments or depletions varied over the time course of infection. Because metabolite exchange drives microbial populations, community composition, and biogeochemical cycling, the significant observed changes in metabolite exudates could appreciably impact these processes during the infection cycle. Therefore, we sought to quantify the chemotactic responses of heterotrophic bacteria as an indication of the potential impact of phage-induced pre-lysis changes in exometabolite composition.

**Fig. 1.**
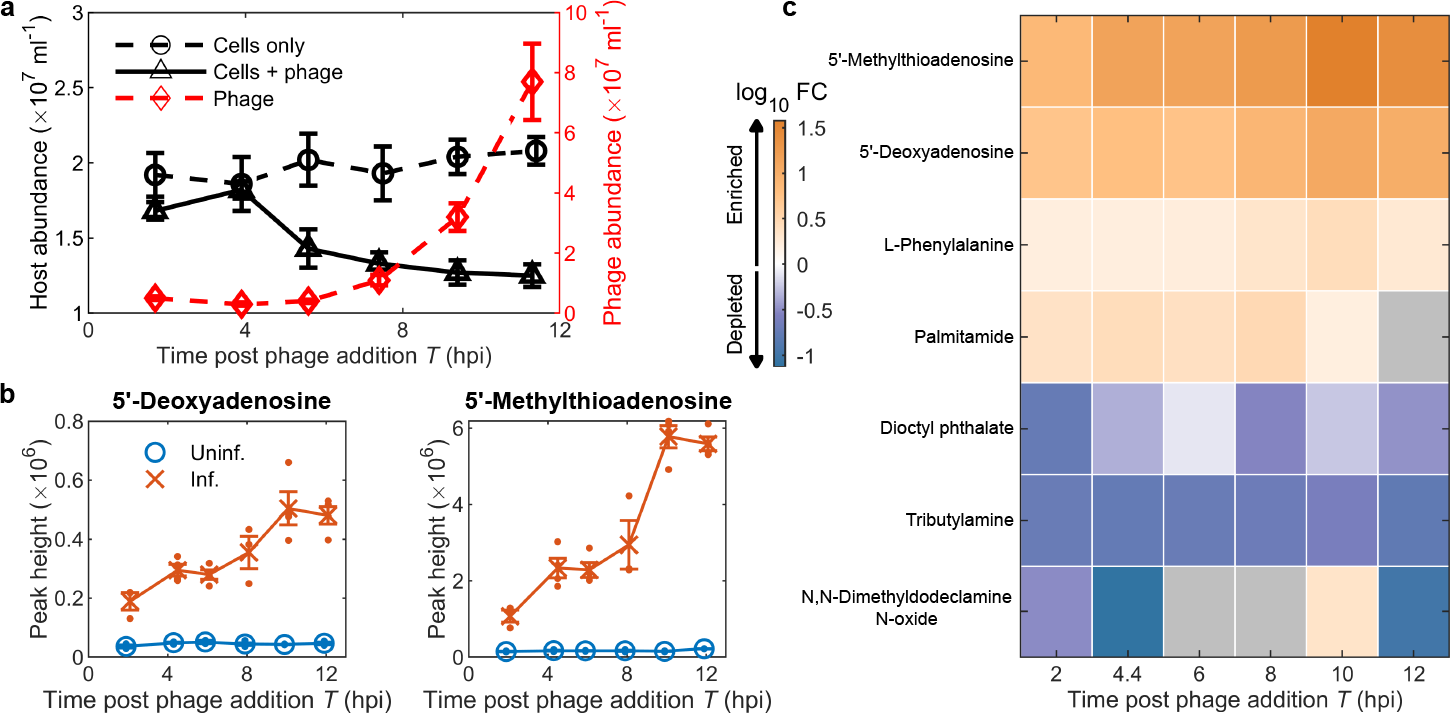
Impact of phage on cyanobacteria population and metabolomic composition. **(a)** Infection dynamics of *Synechococcus* infected with S-SSM5, followed for 12 h post infection (phage addition) (hpi). No significant changes observed in control uninfected populations over the 12 h period; significant (*P <* 0.05) declines in cell populations in phage infected cultures observed at 8,10 and 12 hpi (Extended Data Fig. 1). Adapted from [19]. **(b)** Metabolite peak heights (Methods) for two metabolites significantly enriched during the infection cycle, individual data points shown as dots, error bars indicate standard error (SE). **(c)** Summary of extracellular metabolites in phage infected cultures (Extended Data Fig. 2, Extended Data Table I) exhibiting a high-confidence identification and significant change, shown as log10-fold change (FC) relative to control uninfected cultures, over the course of the experiment. Gray indicates no significant detection (FC *<* 0.1).

### Preferential bacterial chemotaxis towards exudates from infected cells

Having established significant changes in the exudate composition, an extensive panel of microfluidic experiments were conducted to investigate the influence of infection on the chemotactic behaviour of neighbouring heterotrophic bacteria. The model marine bacterium *Vibrio alginolyticus* (Fig. 2a) was chosen due to its well-documented motility and chemotaxis [21–23], and its prevalence in marine microbial communities [24]. The chemotactic response of *V. alginolyticus* to both infected and uninfected exudates was assayed at six time points throughout the infection cycle using an established microfluidic device [20, 25] (Fig. 2b). Briefly, a three inlet device utilises stop-flow diffusion to establish repeatable exometabolite gradients across an observation region (1 mm wide). The three solutions are initially flow-stratified (Fig. 2c; *t* = 0 min), comprising a dilute suspension of *V. alginolyticus* in artificial seawater (Fig. 2c, black dots) flanked on either side by exudates from 0.1 *μ*m filtered infected (Fig. 2bc, red) and uninfected *Synechococcus* cells (Fig. 2bc, white), respectively. The two sets of filtered exudates are sampled from the same time-point in the infection cycle. Upon halting the flow, diffusion establishes infected and uninfected exometabolite concentration gradients across the device (Fig. 2c, red), and time-lapse microscopy measures the chemotactic motility of a *V. alginolyticus* population (≈ 3, 000 cells) over time *t* = 0 − 10 min. An example assay (Fig. 2c) shows the cell response to exudates from the first infection time-point (*T* = 2 hpi). *V. alginolyticus* cells migrate along chemical gradients towards the infected exudates (Fig. 2c, *t* = 1 min) until the gradient dissipates and the bacteria begin to uniformly explore the available space (Fig. 2c, *t* = 10 min). The spatial distribution of cells *P* (*y* | *t*), across the observation channel, *y*, at time, *t*, is shown as a conditional probability (Fig. 2d). The cell population response is compactly characterised by the accumulation index [13, 20] (Fig. 2e), *β*(*t*) = (*n*_*I*_ (*t*) − *n*_*U*_ (*t*))*/*(*N*_*I*_ +*N*_*U*_), where *n*_*I,U*_ (*t*) are the instantaneous number of cells at time *t* within 200 *μ*m of the upper (infected) and lower (uninfected) boundaries (Fig. 2c, white and cyan dashed lines, respectively), and *N*_*I,U*_ are the final cell counts at *t* = 10 min.

**Fig. 2.**
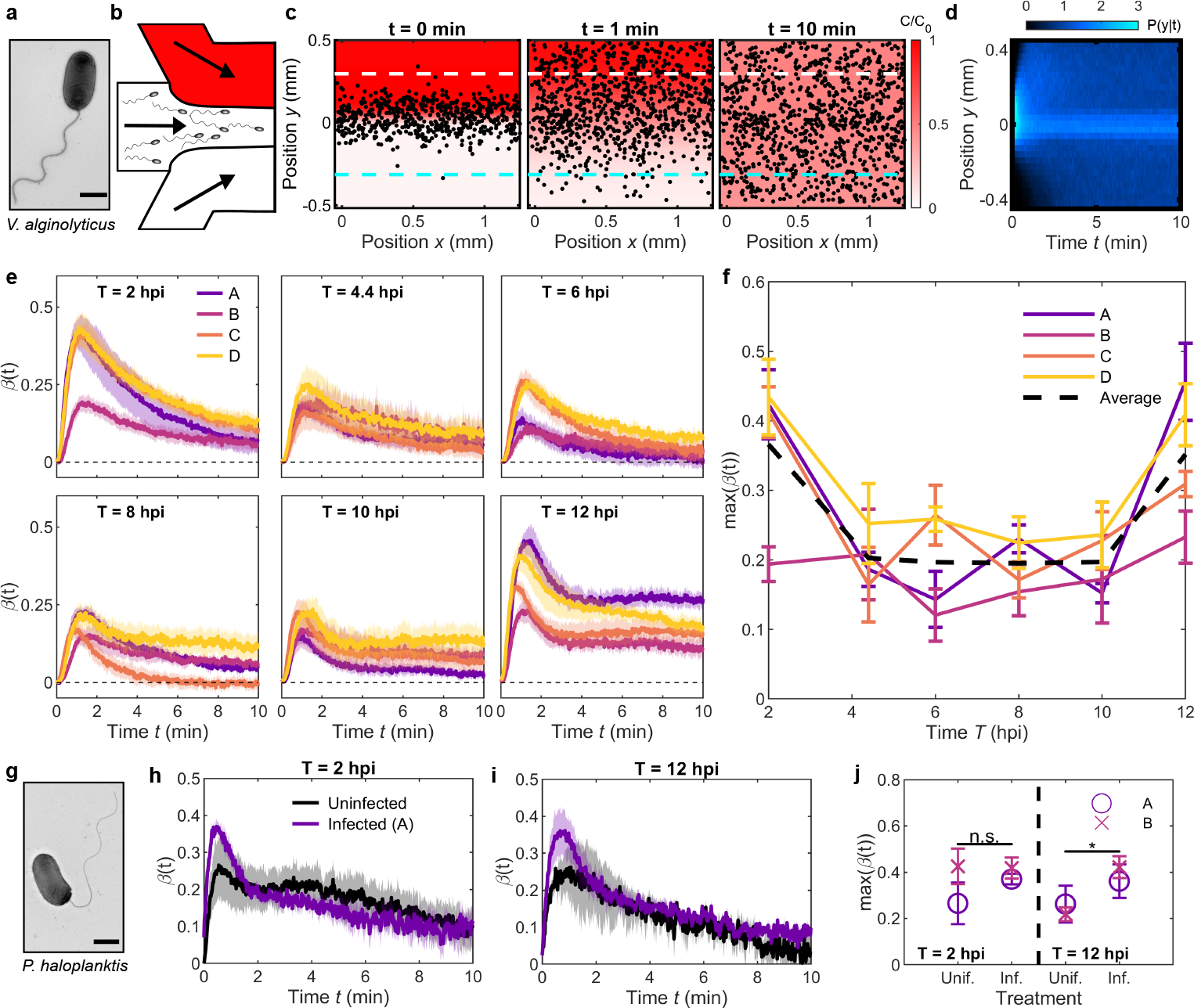
Bacterial chemotaxis towards early-stage infected cyanobacteria. **(a)** TEM image of *V. alginolyticus*. Scale bar 1 *μ*m. **(b)** Three-inlet chemotaxis microfluidic device flows exudates from phage infected cyanobacteria (top, red), cell suspension (*V. alginolyticus*; middle) and exudates from uninfected cyanobacteria (bottom, white) into the observation region (adapted from [20]). **(c)** Measured cell positions (*V. alginolyticus*, black dots; *T* = 2 hpi) at three times *t* after initial flow stratification (*t* = 0 min). Representative gradient measured by fluorescein intensity shown in red. Cells migrate up the infected exudate gradient (*t* = 1 min) followed by dispersal (*t* = 10 min) as the gradient disappears. Accumulation index, *β*(*t*), is measured from cell positions within 200 *μ*m of the top/bottom boundaries (white/cyan dashed lines, respectively). **(d)** Measured bacteria distribution evolves over time and is described by the conditional probability *P* (*y* | *t*). **(e)** Accumulation index, *β*(*t*), for six timepoints post infection for four biological replicates (A-D) (Extended Data Fig. 3). Shading indicates SE. **(f)** Peak accumulation, max(*β*(t)), as a function of time post infection. Strong preferential chemotaxis to the infected cyanobacteria exudates is observed at the end of the infection/lysis cycle, but the strongest and most robust response occurs shortly after infection (*T* = 2 hpi). Error bars indicate SE. **(g)** TEM image of *P. haloplanktis*. Scale bar 1 *μ*m. **(h-i)** *β*(*t*) for *T* = 2 hpi and *T* = 12 hpi, respectively, for *P*.*haloplanktis*. Infected exudates (replicate A, purple) and uninfected exudates (black) challenged against ASW buffer (see also Extended Data Fig. 3 for *V. alginolyticus*). **(j)** Whilst no significant difference is apparent at *T* = 2 hpi for two independent exudate samples (A and B; Extended Data Fig. 3), a statistically significant increase was found for max(*β*(t)) at *T* = 12 hpi (two-tail test, *p* = 0.0454). Error bars indicate SE.

The chemotactic preference of *V. alginolyticus* for exudates from phage-infected (*β >* 0) versus uninfected (*β <* 0) cells were compared at six times during the infection cycle for four biological replicates (independent flasks; Fig. 2, replicates A-D). One biological replicate (Fig. 2ef, replicate B) showed lower phage production, compared to the other three replicates (Fig. 1, Extended Data Fig. 1), suggesting slower infection dynamics and likely reduced impact on exometabolite composition. Across all timepoints, *V. alginolyticus* exhibited preferential chemotaxis towards the infected exudates (Fig. 2ef), exemplified by a strong initial accumulation towards the infected exudate regions with a slow decay towards a uniform cell distribution as chemical gradients naturally dissipate over time via diffusion. The final sampled time point, which is accompanied by frequent lysis events, is characterised by sustained accumulation at long times (Fig. 2e, *T* = 12 hpi). This feature is due to the release of large, slow-diffusing lysate materials, which produce a steadier concentration gradient. In contrast to the expected strong accumulation late in the infection cycle due to the large volume lytic release of organic material [26–28], the strongest accumulation was surprisingly observed early in the infection cycle (Fig. 2e, *T* = 2 hpi). These features are summarised by considering the maximal *β*(*t*) observed at each time point in the infection cycle (Fig. 2f). The chemotactic response is distilled into three stages (Fig. 2f, dashed black line): a sharp initial accumulation during early stages of infection, a decrease to a plateau during the middle to late stages of viral replication, and finally, an accumulation increase following cell lysis.

To explore the ubiquity of preferential bacterial chemotaxis toward phage-infected *Synechococcus* exudates, chemotaxis assays were also performed with a second heterotrophic marine bacterium, *Pseudoalteromonas haloplanktis* (Fig. 2g). Strong chemotaxis to amino acids and cellular exudates have previously been reported for *P. haloplanktis* [29, 30], and the two bacteria are both monotrichous heterotrophs with similar motility patterns [31], common to at least 70% of marine isolates. However, *V. alginolyticus* is known to exhibit higher chemotactic sensitivity to model attractants such as serine in comparison to P. haloplanktis [20]. The chemotactic responses at the first and last infection time points (Fig. 2hi, respectively) were examined independently for the uninfected and infected exudates against artificial seawater (ASW). Similar to *V. alginolyticus, P. haloplanktis* presented a heightened response towards the infected versus uninfected exudates, with a consistent muted response towards the lower-yield infection assay (replicate B; Fig. 2j and Extended Data Fig. 4).

### Marine bacteria are attracted to intact phage-infected cyanobacteria

Whilst relatively long-standing evidence of chemotaxis towards cyanobacteria products exists [30], only recently has experimental evidence established that chemotaxis can enhance exchange between cyanobacteria and heterotrophic bacteria [5]. Such previous experiments – including chemotaxis enhancement via viral infection shown here (Figs. 1 and 2) – typically use a higher concentration of cyanobacteria than naturally-occurring systems in order to produce sufficient quantities of exudates for downstream analysis (e.g. metabolomics). To confirm that the exudation of single phage-infected cyanobacteria is in fact a detectable chemoattractant source for the surrounding bacterial community, we conducted the first microfluidic assays which replaced the exudates (attractant) with a low density of phage-infected or uninfected cells (Fig. 3a). *Synechococcus* were sampled at two points in the infection cycle: *T* = 3 and 7 hpi, respectively (see Methods). The cells were washed and resuspended in ASW, then immediately introduced to the microfluidic device, with ASW used as the control buffer in the third channel. Under these conditions, *Synechococcus* strain WH8102 was experimentally confirmed to be non-motile, and the cyanobacteria remained in the top region of the channel. The number and positions of cyanobacteria present were verified by fluorescence microscopy, with no significant changes detected (Extended Data Fig. 5) throughout the duration of each assay (60 min).

**Fig. 3.**
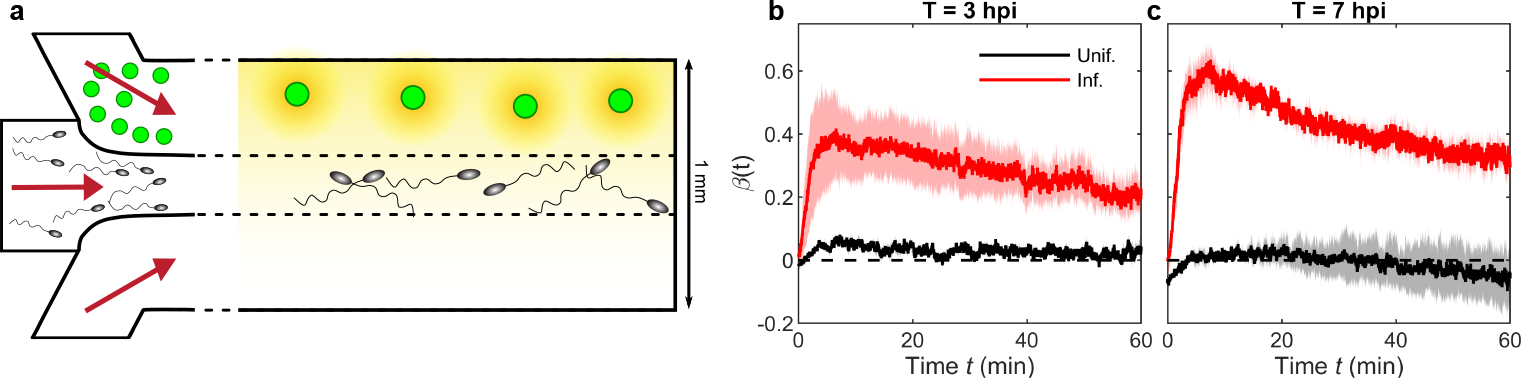
Bacterial chemotaxis toward single, intact, infected cyanobacteria. **(a)** Sketch of the microfluidic assays (not to scale), which use the same three-inlet device as exudate assays (1 mm wide; Fig. 2bc). A dilute intact infected or uninfected *Synechococcus* suspension in ASW was used as the chemostimulus source with *<* 20 cells in the field of view throughout the experiment (Extended Data Fig. 5), a *V. alginolyticus* suspension was injected through the central channel, and ASW was used as the control buffer. Accumulation curves for *V. alginolyticus* **(b)** *T* =3 hpi and **(c)** *T* =7 hpi for uninfected and infected *Synechococcus* (black and red, respectively) over a 60 min period. Shading indicates SE (n=2 and n=3 for uninfected and infected, respectively). At both time points shown, there is significant accumulation towards the phage infected cyanobacteria compared to their uninfected counterparts, which is higher at the later time point in the infection cycle.

**Fig. 4.**
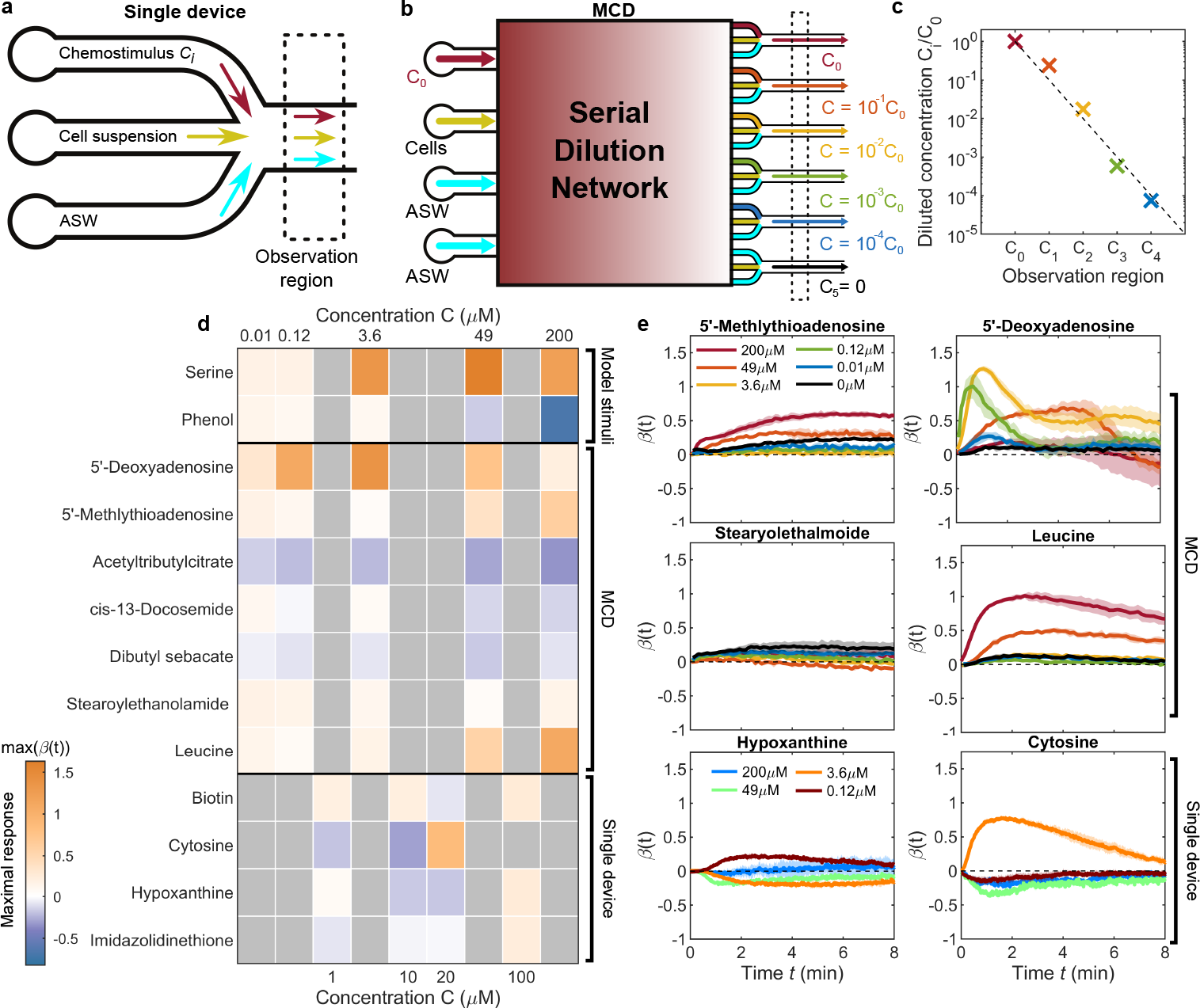
Chemotaxis response of *V. alginolyticus* to identified metabolites. The chemotactic responses of *V. alginolyticus* to identified enriched or depleted metabolites (Fig. 1c) were quantified using **(a)** the single assay device used previously in this study (Fig.2bc, Fig. 3a), and **(b)** a high-throughput multiplexed chemotaxis device (MCD) [20]. The MCD (Extended Data Fig. 6) uses microfluidic serial dilution to automatically dilute and parse out a base chemostimulus solution *C*_0_ to five separate on-chip assays, plus a sixth control condition, for simultaneous chemotaxis screening of different conditions. **(c)** Measured logarithmic dilution of the base chemostimulus (reproduced from [20]). Dashed line corresponds to *C*_*i*_ = *C*_0_ × 10^*− i*^. **(d)** Summary of measured chemotactic response of *V. alginolyticus* taken as the maximum accumulation index during an assay, max(*β*(t)), including a model attractant (serine) and repellent (phenol). Chemoattraction to a range of previously identified enriched or depleted compounds (Fig. 1) was quantified using two different microfluidic platforms (a-b). **(e)** Response curves for a selection of compounds, ranging from strong responses in highly enriched compounds (5’-Deoxyadenosine, 5’-Methylthioadenosine) to null responses in other identified compounds (Stearyolethalmoide). See Extended Data Fig. 7 for full response curves for all identified compounds. Shading indicates SE.

Significant and sustained accumulation was observed towards phage-infected *Synechococcus* (Fig. 3bc, red) at both infection timepoints (*T* =3 and 7 hpi). In contrast, no sustained accumulation by *V. alginolyticus* was observed towards the uninfected *Synechococcus* (Fig. 3bc, black) at either infection timepoint. For uninfected cyanobacteria, numerical experiments recently proposed that chemotaxis increases both the encounter rate and residence time of heterotrophic bacteria with the cyanobacteria phycosphere [5] with interactions occurring on the scale of tens of micrometres. However, in the case of phage-infected cells, significant and sustained bacterial accumulation is observed compared to the absence of a discernible response towards uninfected cells (Fig. 3bc). The continual, modified exudation from the phage-infected cells likely contributes to the sustained accumulation (1 h) over the experiment duration compared to the accumulation timescale (*<* 5 min) observed during the exudate chemotaxis assays (Fig. 2). Since *β*(*t*) is measured within 200 *μ*m of the boundaries, (Methods), the strength of the sustained accumulation suggests an enhanced interaction range in comparison with the theorised diameter of the corresponding phycosphere [3] and moreover, a clear preference for infected cyanobacteria as a nutrient source compared to their uninfected counterparts.

### Bacterial response to individual identified metabolites

From the pool of both polar [19] and non-polar exometabolites (Fig. 1c) identified candidate compounds responsible for changes in bacterial chemotactic behaviour were selected based on both the level of enrichment in phage-infected cultures and ecological relevance. Since the bacterial chemotactic response can change strength or even sign (i.e. from an attractant to a repellent) with compound concentration, the degree of chemoattraction to each candidate compound by *V. alginolyticus* was tested across a wide range of concentrations. Compound screening was initially performed using the chemotaxis device described above (Figs. 2bc, 3a, and 4a). However, given the large number of compounds, concentrations, and replicates required, a rapid screening approach for the candidate compounds was needed. To resolve this issue and achieve high-throughput screening, we employed the recently developed multiplexed chemotaxis device [20] (MCD). This two-layer microfluidic platform utilises serial dilution to enable parallel bacterial chemotaxis assays spanning five orders of magnitude in chemical concentration (Methods; Fig. 4bc and Extended Data Fig. 6). The device receives a base chemical solution of concentration *C*_0_, which is automatically diluted with ASW on a logarithmic scale and parsed to individual observation channels via a microfluidic network (Fig. 4b). The serial dilution process yields five chemical solutions with concentrations *C*_*i*_ = *C*_0_ × 10^*−i*^, × *i* ∈ [0, 4] (plus a sixth control, *C*_5_ = 0), from which the chemotactic responses of *V. alginolyticus* were simultaneously measured. The six individual on-chip assays were designed to have identical channel geometry and flow conditions as the single chemotaxis device (Fig. 2bc), and the performance of chemotaxis screening was previously verified between the MCD and single assay device [20].

The observed peak bacterial chemotaxis response summarises the results of this panel of assays (Fig. 4d), reported by the maximum value of the accumulation index *β*(*t*) (or minimum in the case of chemorepellents; see also full response curves in Extended Data Fig. 7). Some metabolites were tested with the single assay device (*C* = 1 − 100 *μ*M; Fig. 4d, below dashed line), but the majority were quantified using the MCD (*C* = 0.012 − 200 *μ*M; Fig. 4d, above dashed line). Serine and phenol were included as positive and negative controls, respectively. *V. alginolyticus* is well-known to respond to serine concentrations above 0.2 *μ*M for this model attractant [20, 22], while high concentrations of phenol act as a repellent [20, 32]. For the identified metabolites, significant positive responses were also measured in specific initial concentrations, including leucine (49 *μ*M, 200 *μ*M), cytosine (20 *μ*M), 5’-methlythiadenosine (200 *μ*M) and across almost all tested concentrations of 5’-deoxyadenosine (Fig. 4e). Other enriched compounds, such as stearoylethanolamide and hypoxanthine, did not elicit a measurable response from *V. alginolyticus* across the range of examined concentrations.

## DISCUSSION

In this study, microfluidics and metabolomics were incorporated to directly quantify the chemotactic motility of heterotrophic bacteria in response to phage infected cyanobacteria and their exometabolites. During phage infection, host cells become metabolically distinct from their uninfected counterparts. This observation suggests time-dependent changes in the bulk dissolved metabolite pools and phycosphere chemical composition between infected and uninfected cells as infection progresses. Recent work [19] in the same host-virus system (Fig. 1a) demonstrated significant changes in the polar metabolite composition (largely enriched) of infected *Synechococcus* and the enhanced release of various compounds during middle to late infection, including nucleotides, amino acids, vitamins, and signalling molecules. However, non-polar metabolites – which often comprise secondary metabolites that are non-essential for survival – can provide advantages to organisms under different environmental conditions [33]. Here, we quantified non-polar metabolite changes (Fig. 1c) over the course of viral infection. While cyanophage control of host nucleotide biosynthesis, photosynthesis, and nutrient acquisition is well documented [19, 34–38], viral alteration of secondary metabolite pathways remains far less explored. Recent work has highlighted the role of lysogenic viral infection in altering secondary metabolite synthesis and release [39], important as the commercial growth of cyanobacteria for secondary metabolite production is common [40–42]. Our present work further reveals a diversity of non-polar exometabolite changes over the course of a lytic cyanophage infection, identifying 7 compounds significantly different from uninfected control cultures (Fig. 1c, Extended Data Table I).

Infected *Synechococcus* exhibited increased release of ribonucleosides, amino acids and fatty amides, and depletion of benzenoids, amines, and amine oxides. In particular, the two most elevated compounds throughout the infection cycle – the primary metabolites 5’-Deoxyadenosine and 5-Methylthioadenosine (MTA) – elicited strong positive chemotactic responses in *V. alginolyticus* when independently tested (Fig. 1c and Fig. 4de). Dissolved MTA has been found in surface waters of the North Atlantic Ocean in an abundance positively correlated with chlorophyll *a* and pico/nano-eukayrotes [39]. Whilst no correlation has been found between MTA and *Synechococcus* abundance, MTA could be an indicator of active or recent infection of natural *Synechococcus* populations, but remains to be confirmed. Two of the identified compounds (tributylamine and N,N-Dimethyldocylamine) are known or suspected antimicrobial agents [43, 44], both of which were depleted at both 2 hpi and 12 hpi, corresponding to the strong positive chemotactic responses of *V. alginolyticus* (Fig. 1c, Fig. 2ef). Depletion in antimicrobial agents could elicit and/or enhance positive chemotactic response from nearby bacteria or protists, an intriguing connection that warrants further study. Our identifications and measured phage-induced changes are consistent with previously reported examples for other cyanobacteria-phage systems, for example the enrichment of MTA and depletion of phthalates [45–48], implying our results are not specific to our chosen host (*Synechococcus* WH8102) and phage (S-SSM5) strains, but rather might be generalisable to wider cyanobacteria-phage-bacteria interactions. Taken together, this plethora of phage-induced exometabolite changes suggests significant potential for viral infection to augment the chemotactic behaviour of heterotrophic bacteria throughout the infection cycle.

Chemotaxis towards large lysed diatom cells (≈ 200 *μ*m) was previously reported [12] to act as a strong yet ephemeral (e.g. *<* 10 min) nutrient source for the surrounding microbial community through the sudden release of dissolved organic matter (DOM). Mathematical modelling based on these data suggested that viral lysis of plankton would be insufficient to sustain motile heterotrophic bacterial populations unless under conditions of high cell density, such as plankton bloom scenarios. Through an extensive panel of microfluidic assays (Fig. 2bc), we showed here that metabolite exudates, released during the early stages of viral infection (Fig. 2ef), are much stronger chemoattractants to heterotrophic marine bacteria (e.g. *V. alginolyticus* and *P. haloplanktis*; Fig. 2fhi) compared to exudates from uninfected cells. At all stages of infection, there is consistent chemotactic preference towards exudates derived from phage-infected host cells. There is an expected strong response near the time of lysis, but a second response of equal magnitude is also observed early in the infection cycle (Fig. 2f). These results are further supported by replacement of exudates with live-infected cyanobacteria (Fig. 3), where heterotrophic bacteria maintain a significant accumulation towards infected cells and importantly do not display the same response to uninfected cells. These results also form the first direct experimental evidence of heterotrophic bacteria chemotaxing towards live, infected cyanobacteria. Recent work [5] has shown that chemotaxis enhances exchange between cyanobacteria and heterotrophic bacteria, with numerical experiments suggesting that short-range interactions are responsible. Hence, the rate of exudation and/or composition of exuded chemoattractants from uninfected cells might be insufficient to establish the necessary chemical gradients to elicit the population-scale accumulations observed here towards infected cyanobacteria.

The prevalence of virus-infected microbes [9] combined with their influence on the spatio-temporal distribution of chemotactic heterotrophic bacteria illustrated here has significant implications for marine microbial ecology. Inter-microbe distances in the open ocean are generally believed to be on the order of 100^*′*^s of microns [49], making for sparse nutrient sources. Previously it has been assumed that the range of the detectable phycosphere produced by picoplankton such as *Synechococcus* is at most on the scale of a few tens of microns [3, 5]. Similarly, ephemeral nutrient patches released during cell lysis typically persist on the scale of a few minutes [12]. Thus, heterotrophic bacteria have an extremely limited target size and time window for random encounters with small phycospheres and plumes from lysed cells, respectively. Given these relatively short interaction time scales, nutrient uptake from these sources may provide a comparatively more limited advantage relative to infected cyanobacteria. The latter is a potentially more stable nutrient source: Firstly, the chemotactic interaction range with phage-infected cells is greatly enhanced compared to their uninfected counterparts (Fig. 3bc). Secondly, phage-infected cells function as a long-lasting (e.g. hours) nutrient source compared to the ephemeral nature of lysis events (e.g. minutes). Finally, as shown by enrichment of secondary metabolites, these biological hotspots enable chemotactic heterotrophic bacteria faster access to a wider and/or enriched range of compounds (Fig. 4) as compared to non-chemotactic bacteria, which potentially shapes community composition and provide additional access to important sources of nitrogen and sulphur. Overall, our work demonstrates a new viable mechanism for nutrient recycling that could influence our understanding of the transfer of carbon and other nutrients at the trophic scale.

## METHODS

### Cell culturing

*Vibrio alginolyticus* (YM4; wild-type) from − 80°C stock were grown overnight in Marine 2216 media (Difco) by incubating at 30°C and shaking at 600 revolutions per minute (RPM). The overnight culture was diluted 100-fold into fresh pre-warmed 2216 media and grown for three hours (30°C, shaking at 600 RPM) to O.D. 0.2. 2 ml of culture was then centrifuged (1,500 relative centrifugal force (RCF) for 5 min), washed twice with artificial seawater (ASW), and resuspended in 0.5 ml of ASW. ASW was prepared according to the NCMA ESAW Medium recipe with some modifications [50, 51].

*Pseudoalteromonas haloplanktis* (ATCC 700530) from −80°C stock were grown in 2216 media for 24 hours at room temperature without agitation. The overnight culture was then centrifuged (1500 RCF for 5 min), and 4 ml of culture was washed twice and resuspended in 0.5 ml of ASW.

For the chemotaxis of *V. alginolyticus* to live cyanobacterial cell assays, axenic *Synechococcus* WH8102 (CCMP 2370) were grown in SN media [50, 51], prepared with ASW, in sterile 40 ml polystyrene culture flasks at 22°C on a 14 h : 10 h light-dark cycle at 50 *μ*mol photons m^*−*2^ s^*−*1^. Culture growth was tracked using a SpectraMax ID3 plate reader (Molecular Devices) and cell counts were measured using a CytoFLEX flow cytometer (Beckman Coulter).

### *Synechococcus* exudate generation and collection

Growth and infection conditions were described previously [19]. Briefly, axenic *Synechococcus* strain WH8102 (CCMP2370) was grown in SN media prepared with sterile 0.1 *μ*m filtered, autoclaved natural seawater. Cultures were maintained at 20°C on a light-: dark cycle of 14 h : 10 h at 60 *μ*mol photons m^*−*2^ s^*−*1^. Infection with the lytic T4-like myovirus S-SSM5 was initiated at the onset of the light cycle with a MOI of 1.1. Four biological replicates were prepared, using independent flasks, for both phage-infected and control uninfected cultures. Phage titer was determined by the most probable number (MPN) assay [52, 53] using three independent dilution series on axenic *Synechococcus* WH8102 grown in 96 well plates under growth conditions described above, and phage stocks were used within 14 days of generation. After 1 h, cells were centrifuged, supernatant was removed, and cells were resuspended in spent culture medium. Host and phage concentrations were monitored via flow cytometry as described previously [19]. *Synechococcus* starting concentrations were 1.9*×*10^7^*±*0.3*×*10^7^ and 1.7 × 10^7^ ± 0.1× 10^7^ cells ml^*−*1^ in control uninfected and phage infected cultures, respectively. Samples were collected between 2 and 12 h post-initial phage addition every 2 h during the daylight portion of the light : dark cycle, by gently mixing and pouring 100 mL into a sterile 250 mL polycarbonate flask. Cell and phage abundance samples were preserved with a final concentration of 0.1% EM grade glutaraldehyde (Acros Organics), stored at 4°C for 15-30 min, then flash-frozen in liquid nitrogen and stored at 80°C until counted via flow cytometry. For exometabolite and chemotaxis assay filtrate samples, ≈ 2 × 10^8^ *Synechococcus* cells were filtered onto 0.11 *μ*m pore size PES filters (Sterlitech), and the filtrate was stored at − 20°C until analysis. All downstream analyses were performed with a minimum of three biological replicates. Further details are described elsewhere [19].

### Metabolomics

20 ml of filtrates were lyophilised until dry, then resuspended in 1 ml of ice cold methanol, vortexed twice for ten seconds and then bath sonicated in ice water for 10 min before centrifugation (10,000 RCF for 5 min at 10°C). Supernatants were collected into a new tube and dried using a vacuum concentrator (Thermo Speed-Vac). Dried extracts were resuspended in 150 *μ*l ice cold methanol containing IS (Tables S1-2). Vortexing, sonication, and centrifugation were repeated, with supernatants then filtered using 0.22 *μ*m PVDF microcentrifuge tubes and subsequent filtrates collected in glass vials for LCMS analysis. Metabolites were analysed using reverse phase LC-MS/MS; parameters are described in the Supplemental Material(Tables S1-2).

### Compound identification

Feature detection was performed with MZmine 2.39 using the parameters defined in the Supplementary Materials (Table S3 and associated .xml batch file). Compound identification was performed using the Global Natural Product Social Molecular Networking (GNPS) database [54]. Identifications were required to have at least 4 matching ions and a cosine score greater than 0.7. Final target compounds were identified from the filtered list by the following seven criteria: (i) greater than 1 min retention time to exclude not-retained features, (ii) compulsory tandem mass spectra, (iii) a signal-to-background ratio of at least three in at least one sample, (iv) minimum average peak height in at least one sample type greater than 100,000 counts, (v) minimum range between at least two sample types greater than two, (vi) modelled as a Student’s t-distribution with the 95% confidence interval (lower bound) greater than 10,000 counts, and finally (vii) tested for significance with a two-way ANOVA considering the time and treatment as independent and interaction effects and removed features not significantly different for the phage treatment vs the control. The resulting features and their best matching identification are shown in Fig. 1c, Extended Data Fig. 2 and Extended Data Table I.

### Microfluidic chemotaxis assays

Microfluidic devices were fabricated through standard soft lithography techniques [55]. Photolithography was used to make photoresist (Kayaku Advanced Materias, formally Microchem) molds on a silicon wafer Microfluidic channels were cast with polydimethylsiloxane (PDMS; Dow Corning SYLGARD 184) and plasma bonded to standard glass microscope slides. Prior to use, channels were pre-treated by first flowing a 0.5% (w/v) bovine serum albumin solution (BSA; Sigma Aldrich) to reduce cell adhesion to the channel surfaces, then flushed with ASW. The three microchannel inlets (Fig. 2b) carried the chemostimulus, bacterial cells (*V. alginolyticus* or *P. haloplanktis*) suspended in ASW, and control solutions, respectively. For the exudate chemotaxis experiments (Fig. 2), the chemostimulus/control solutions were the infected/uninfected *Synechococcus* exudates, respectively. For the identified compound experiments (Fig. 4), the chemostimulus/control solutions were the identified compounds resuspended in ASW, and ASW, respectively. The three solutions were flow stratified for a minimum of 2 min using a syringe pump (Harvard Apparatus) with flow rates controlled by syringe size to maintain stream width ratios of 4:1:4. Imaging of the exudate experiments (Fig. 2c) was performed using phase-contrast microscopy (Nikon Ti-E; 4 ×, 0.13 NA objective) at 1 fps over the course of 10 min using a CMOS camera (Blackfly S; Teledyne FLIR). For each infection replicate (A-D; Fig. 2e and Extended Data Fig. 3), three technical replicates were collected by restarting the flow, then fully repeated three times with fresh cell suspensions from different cultures for a total of nine replicates.

### Cell identification and analysis

Cell locations (Fig. 2c) were determined to sub-pixel accuracy using available image analysis tools [56] in MATLAB (2021a, MathWorks). The background of each image was removed via mean image subtraction, and a bandpass filter applied to further enhanced cell contrast. Cells were identified through intensity thresholding of the filtered images, and cell locations were determined from the centroid of each identified object. The outcome of this analysis was the spatial distribution of cells as a function of time for each assay (Fig. 2d). Subsequently, the accumulation index *β*(*t*) was determined (Fig. 2e) from cells within 200 *μ*m of the microchannel side walls as described in the main text.

### Whole-cell chemotaxis assays and phage infection conditions

To directly determine the chemotactic affinity of heterotrophic bacteria to intact, infected cyanobacteria, six microfluidic experiments were conducted (Fig. 3) each on different days: three with phage infection and three control, uninfected experiments. Both uninfected and phage-infected experiments were performed identically, with the substitution of phage addition for an equivalent volume of SN media in the uninfected experiments. All infection experiments were performed within a week using the same phage stock, with the infectious phage titer of the stock determined to be 1.4 × 10^8^ pfu ml^*−*1^ via plaque assay method with *Synechococcus* WH8102 as the host. One day prior to experiments, exponential phase *Synechococcus* cultures were diluted with fresh SN media to a cell concentration of 7–9 × 10^5^ cells ml^*−*1^. On the day of experiments, at the onset of the light cycle, culture concentrations were adjusted to 1 × 10^6^ cells ml^*−*1^ as necessary with fresh SN media. A volume of phage stock (or SN media for uninfected controls) was added to achieve a target multiplicity of infection (MOI) of 3. Cultures were then mixed for one min every five min for 45 min in 50 ml polypropylene conical tubes, then centrifuged at 2,000 RCF for 15 min at 20°C. Supernatants were decanted and cells were resuspended in an equal volume of fresh ASW and transferred to sterile 50 ml polystyrene tubes. At approximately two and six hours post resuspension (*T* = 3 and *T* = 7 hpi), live cell concentration was assessed by flow cytometry and 10 ml of culture was collected for chemotaxis assays. Sub-samples were fixed in 0.5% glutaraldehyde for cell and phage quantification [57, 58].

For the live-cell chemotaxis assays (Fig. 3), 10 ml of cells were centrifuged (2,000 RCF for 15 min) and washed twice. Cells were resuspended in 1 ml of ASW and adjusted as necessary with ASW to a final concentration of 3–4 × 10^5^ cells ml^*−*1^. The final cell suspension was loaded into a 1 ml gas tight syringe (Hamilton; 1000 series). Microfluidic chemotaxis assays were conducted as described above with the live-cell solution (infected or uninfected) as the chemostimulus, *V. alginolyticus* suspension (10^9^ cells ml^*−*1^ in ASW) in the centre, and ASW as the control solution. Images were collected for 60 min post flow stratification (Zeiss Axio Observer; 10 ×, 0.25 NA objective) using a CMOS camera (Blackfly S; Teledyne FLIR) at 1 FPS. Fluorescent images were collected immediately prior to and after the one hour chemotaxis assay to count the number of *Synechococcus* present, with pairwise t-tests confirming no statistical changes of the number of *Synechococcus* in the observation region throughout each assay (Extended Data Fig. 5).

### Multiplexed chemotaxis device

A brief summary of the multiplexed chemotaxis device (MCD) is provided below, whereas the full details of the fabrication, characterization, and operation may be found in previous work [20]. The MCD performs six simultaneous chemotaxis assays on a single two-layer microfluidic chip (Fig. 4b, Extended Data Fig. 6), each of which are identical to the standard three-inlet chemotaxis assay described above. Serial dilution is used to produce five logarithmically diluted chemostimulus solutions from a base concentration *C*_0_ (Fig.4bc), by mixing with a given buffer solution. The sixth channel performs a control assay with no chemostimulus present (i.e. *C*_5_ = 0). The observation channels were designed fto have the same flow rates and geometry as the individual chemotaxis assays used for the exudate experiments (Fig. 2). The MCD was fabricated using the same soft lithography methods described above. For this two layer device, the dilution layer was first bonded to a glass substrate, and subsequently, the cell injection layer was aligned and bonded to the dilution layer. The MCD was pre-treated a 0.5% w/v BSA, and fluid flow was driven by a single pressure controller (Elveflow OB1). Pressures of 200 mbar and 140 mbar were used for the dilution and cell injection layers, respectively. Between each assay, the fluid inputs were flowed for a minimum of 2 min to achieve the necessary fluid stratification. Spatio-temporal cell distributions were determined by imaging the cells using phase-contrast microscopy (10×, 0.3 NA objective; Nikon Ti-E) with a sCMOS camera (Zyla 5.5, Andor Technology). An automated computer-controlled stage rapidly imaged each channel sequentially with an effective imaging period of 8 s over the course of approximately 10 min.

### TEM imaging

Initial cultures of *V. alginolyticus* and *P. haloplanktis* were grown as previously described before the following final suspensions were prepared: (i) 4 ml of *V. alginolyticus* culture was washed and resuspended (1,500 RCF for 5 min) in 1 ml of fresh 2216 media, and (ii) 1 ml of *P. haloplanktis* culture was washed and resuspended (1,500 RCF for 5 min) in 1 ml of fresh 2216 media, then diluted 10 times in double distilled water (DDW). For both species, 4 *μ*l of cell suspension was applied to a glow discharged copper mesh carbon coated grid and allowed to adsorb to the grid for 30 s. The grid was briefly washed in DDW followed by staining with 1% Aqueous Uranyl Acetate, and allowed to dry fully before imaging. The grid was imaged using a FEI Morgagni transmission electron microscope (FEI; Hillsboro, OR) operating at 80 kV and equipped with a CMOS camera (Nanosprint5, AMT).

## Supporting information

Batch files containing feature detection parameters

## DATA AND CODE AVAILABILITY

Data files and code used during the current study is publicly available at BCO-DMO [59, 60].

## ACKNOWLEDGEMENTS

We thank J.B. Raina for helpful discussions. TEM samples were prepared and imaged by the Brandeis Electron Microscope Facility.

## AUTHOR CONTRIBUTIONS

R.J.H., J.M., T.N., J.S.G, and S.A.F. designed the research; S.A.F. conducted the culture experiments for exudate generation; R.J.H. performed exudate and target compound chemotaxis assays and analysed corresponding data; S.K. analysed the exometabolomics samples; S.K., B.B., T.N., J.M., and S.A.F. analysed the exometabolomics data; J.M. performed *P. haloplanktis* and whole-cell chemotaxis assays; R.J.H. and J.M. analysed the corresponding data; R.J.H. and M.R.S. contributed microfluidic tools and TEM imaging; R.J.H., J.M., J.S.G., and S.A.F. discussed the results and wrote the manuscript with contributions from all authors.

---

**Extended Data Fig. 1.**
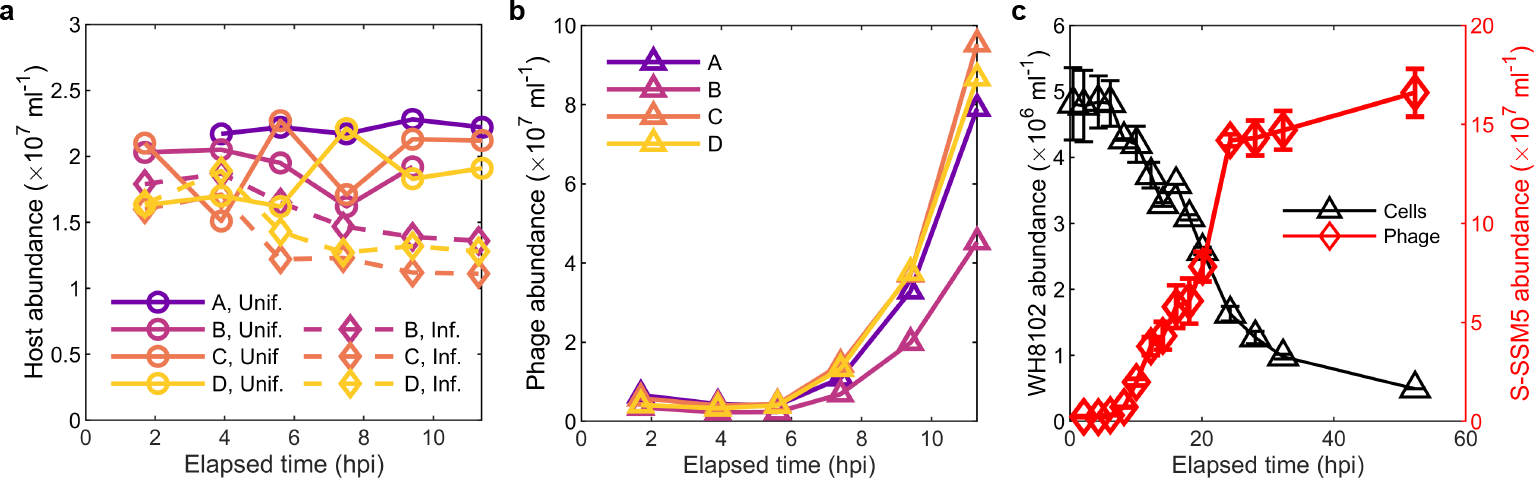
Replicate infection dynamics of *Synechoccocus* WH8102 infected with S-SSM5. Adapted from [19]. **(a)** Host cell abundance in both the uninfected and infected treatments over the 12 h experimental period for four replicates (A,B,C,D). **(b)** Phage abundance in the infected treatment for the four replicates, where the B treatment displayed significantly reduced phage production compared to the remaining three treatments. **(c)** General infection dynamics, followed over 55 h post phage addition, capturing multiple infection cycles. See also Fig. 1.

**Extended Data Fig. 2.**
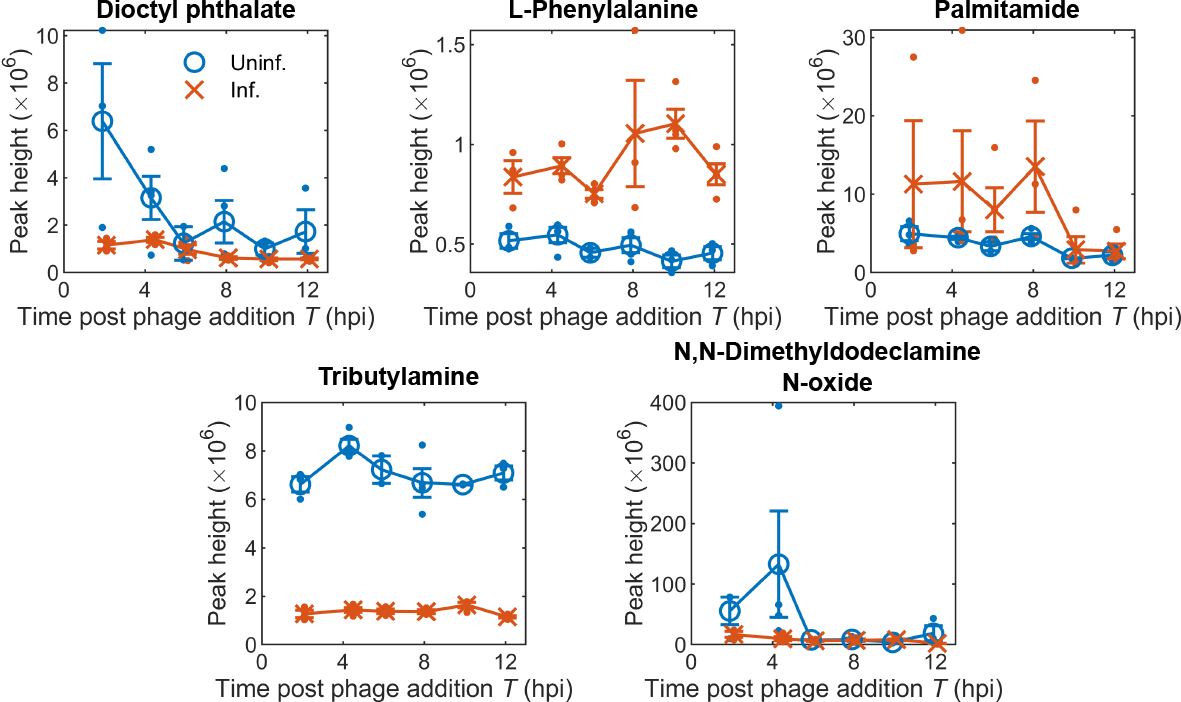
Time-dependent metabolite peak heights. Summary of peak heights from detected and identified extracellular metabolites (Fig. 1c) exhibiting a high-confidence significant difference between phage infected and uninfected treatments. Dots indicate single data points, errorbars denote SE.

**Extended Data Fig. 3.**
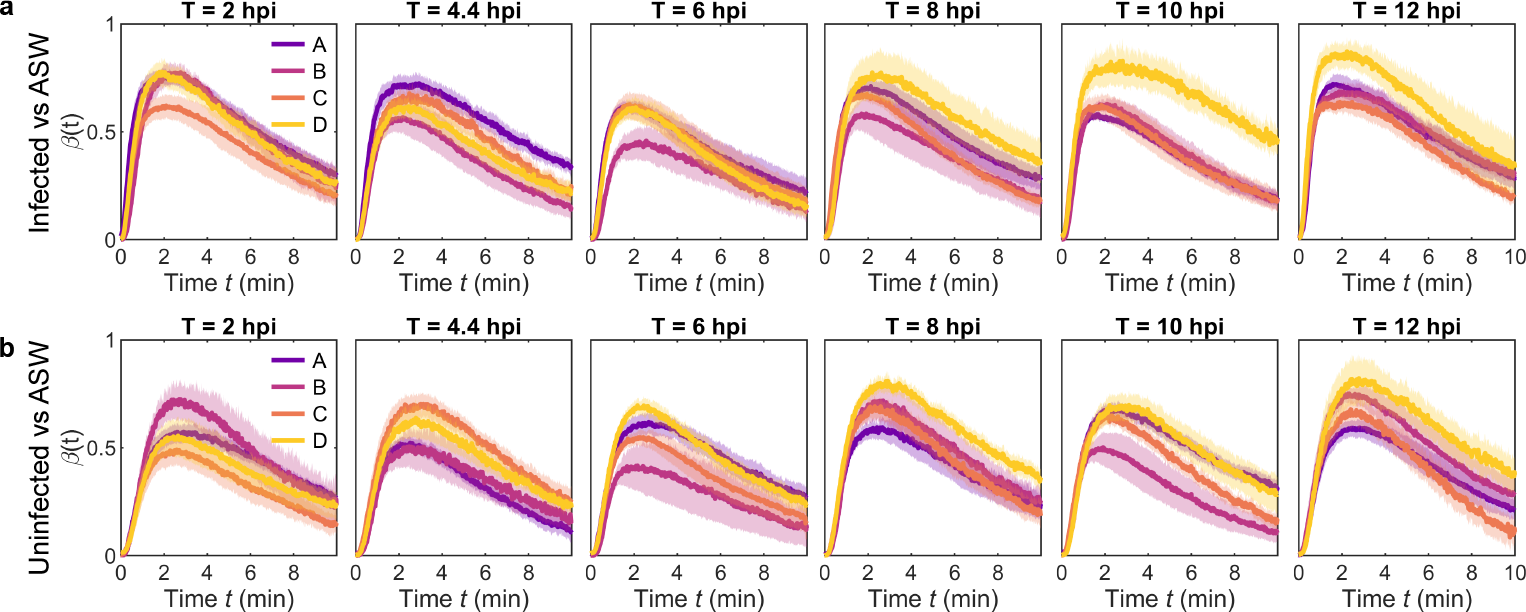
Chemotaxis of *V. alginolyticus* to exudates from phage infected and uninfected cells relative to ASW. Accumulation index *β*(*t*) for chemotaxis of *V. alginolyticus* between **(a)** infected and **(b)** uninfected cyanobacteria exudates when screened against ASW. Assays were performed using the three-inlet single assay device (Fig. 2b) for the six sampled timepoints during the infection cycle. A consistently strong, positive chemotactic response is observed in all cases, with no significant difference in the independent responses to either the infected or uninfected exudates (see Fig. 2e-f). Shading indicates SE.

**Extended Data Fig. 4.**
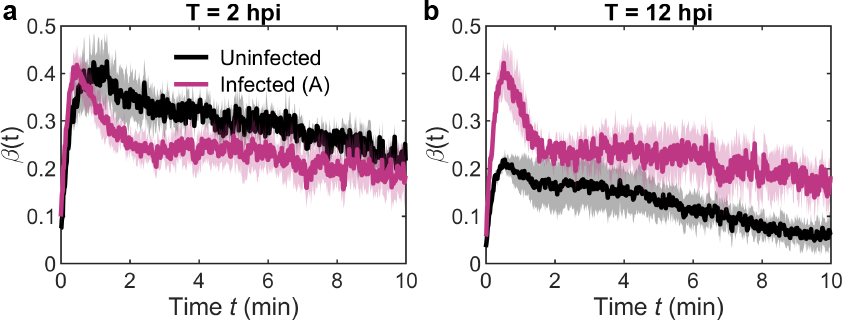
Chemotaxis of *P. haloplanktis* to infected and uninfected exudates (B replicate). **(a)** Similar to *V. alginolyticus* (Fig. 2f), a weaker response was observed in the (B) infected exudate sample, with **(b)** a clear increase in response at the end of the infection cycle (see Fig. 2h-j). Shading indicates SE.

**Extended Data Fig. 5.**
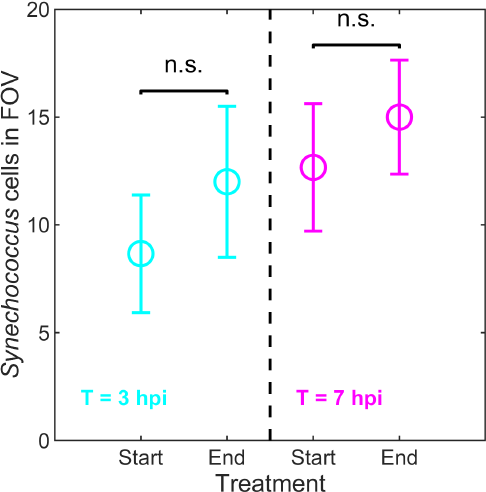
Number of *Synechococcus* present during chemotaxis experiments with live phage infected cells as the chemostimulus. Fluorescent images were collected immediately prior to and after the one hour chemotaxis assays (Fig. 3). The number of *Synechococcus* in the field of view (FOV) were counted, and pairwise t-tests confirmed no statistically significant changes in the number of infected cells in the observation region throughout each assay. Errorbars indicate SE.

**Extended Data Fig. 6.**
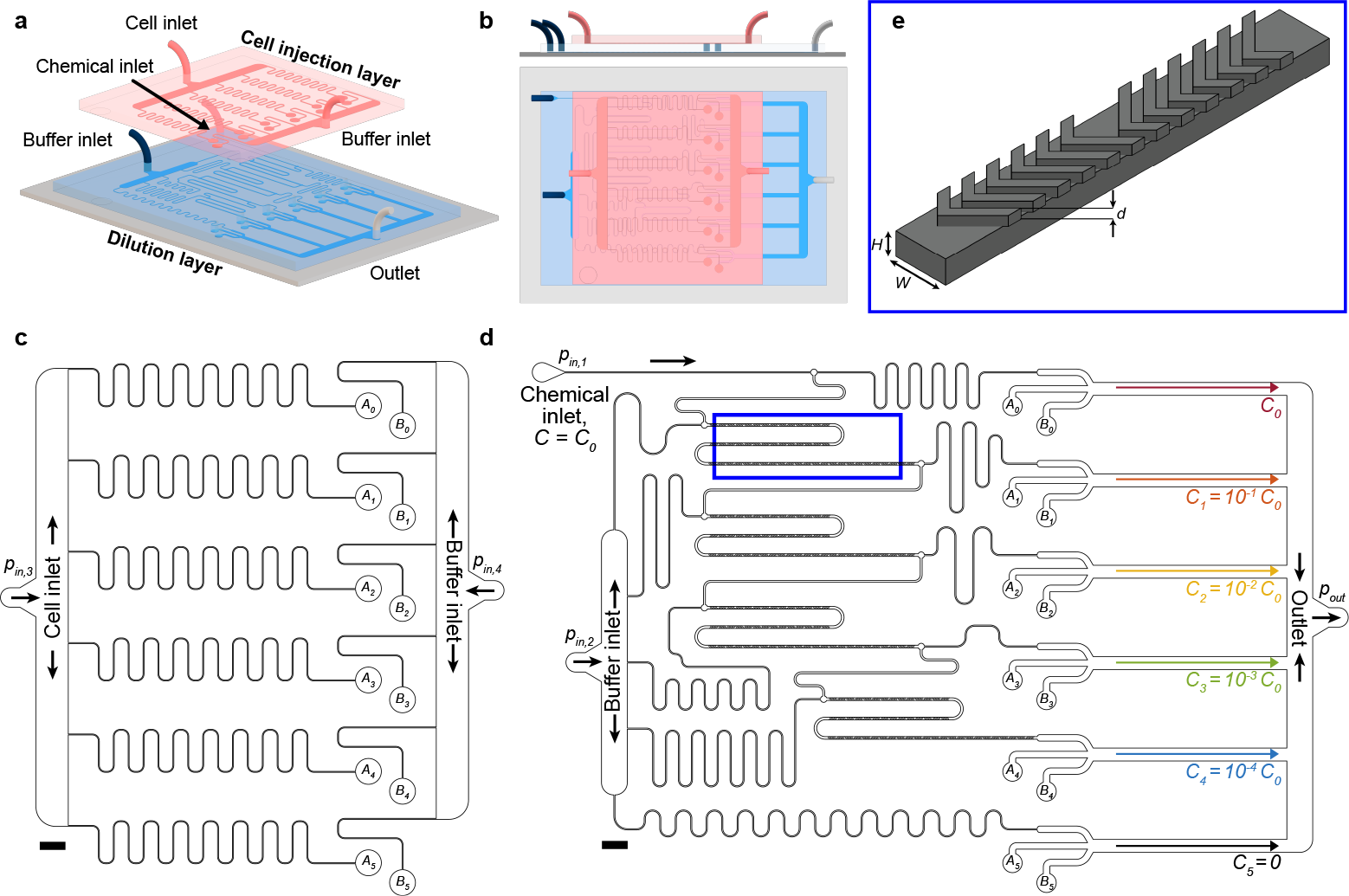
Schematic of the multiplexed chemotaxis device (MCD). **(a,b)** The MCD is a two-layer microfluidic platform designed for rapid screening of bacterial chemotaxis. The device receives four fluid inputs: two buffer solutions (ASW), one cell suspension (*V. alginolyticus* or *P. haloplanktis*), and one chemostimulus of base concentration *C*_0_. The **(c)** cell injection layer (red in a and b) supplies several buffer streams and cell suspension streams via vertical ports (*A*_*i*_, *B*_*i*_) to the **(d)** dilution layer, where six separate on-chip chemotaxis assays are performed simultaneously. The dilution layer uses serial dilution to produce five logarithmically diluted chemostimulus solutions (*C*_*i*_ = *C*_0_ × 10^*− i*^, *i*∈ [0, 4]) and a control channel (*C*_5_ = 0) totalling six chemotaxis assays. **(e)** The device utilises herringbone mixing channels (blue box in d) to ensure accurate chemostimulus dilution. See also Fig. 4; panels are adapted from [20].

**Extended Data Fig. 7.**
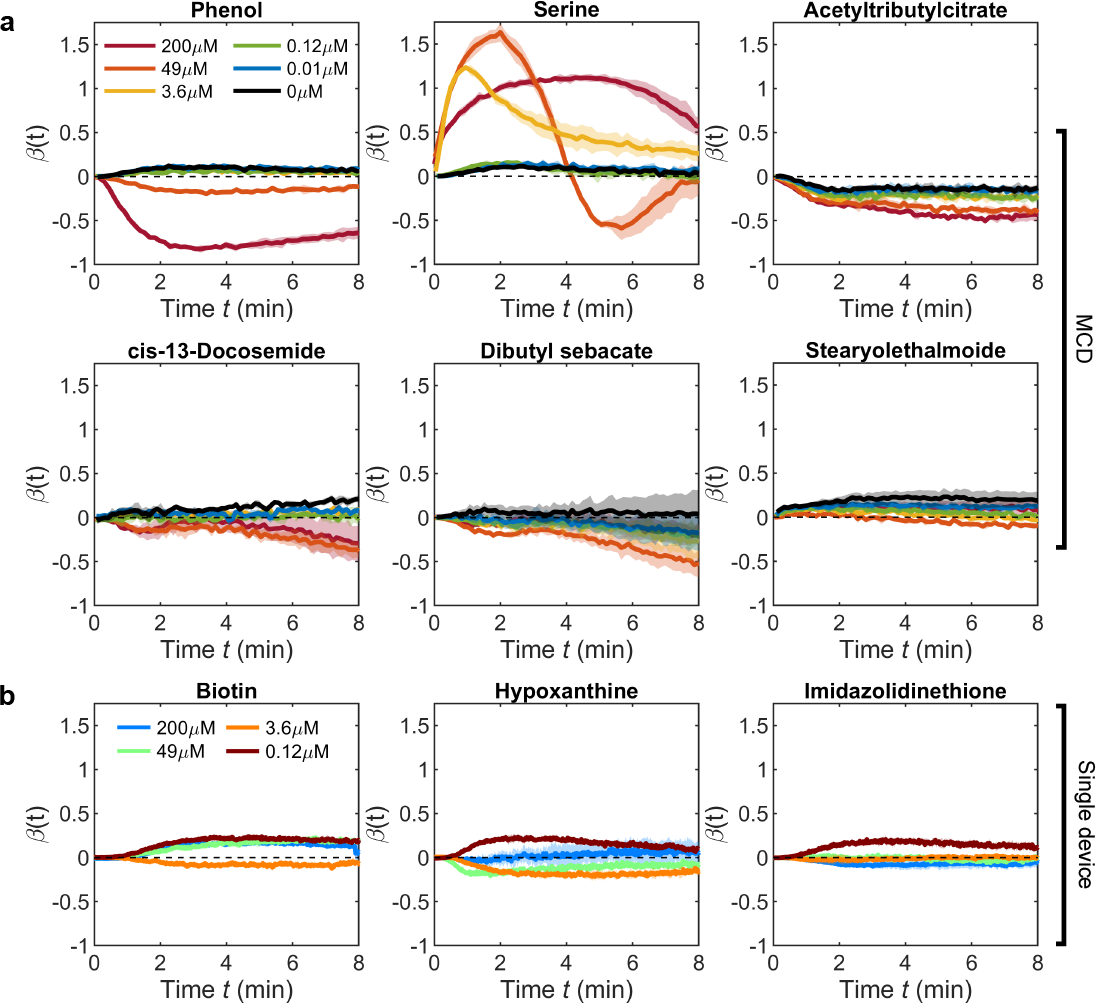
Chemotactic responses to identified compounds. Response of *V. alginolyticus* to identified metabolites (Fig. 4d) using **(a)** the MCD (Fig. 4b) and **(b)** the single assay device (Fig. 2bc).

**Extended Data Table I:**
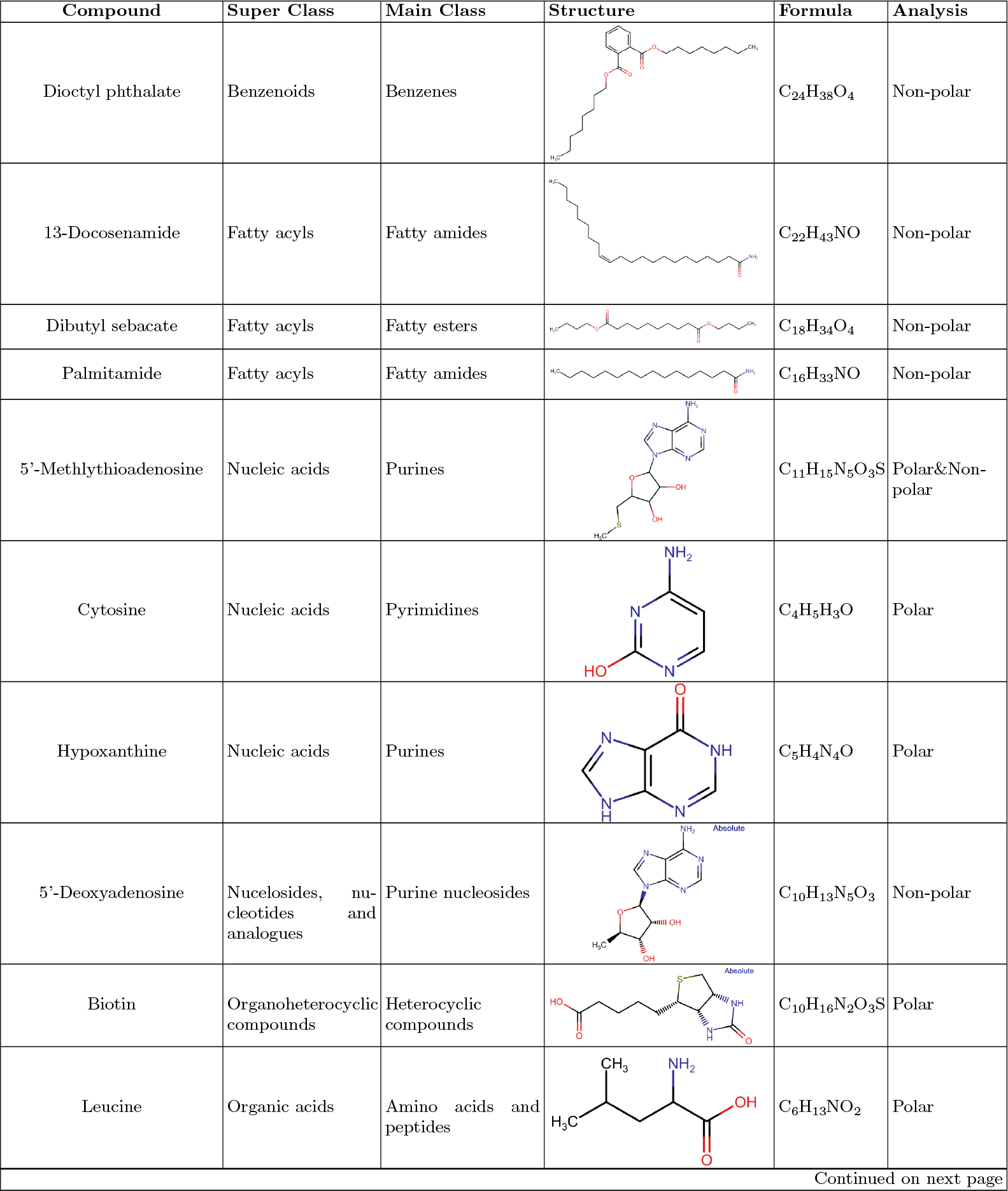
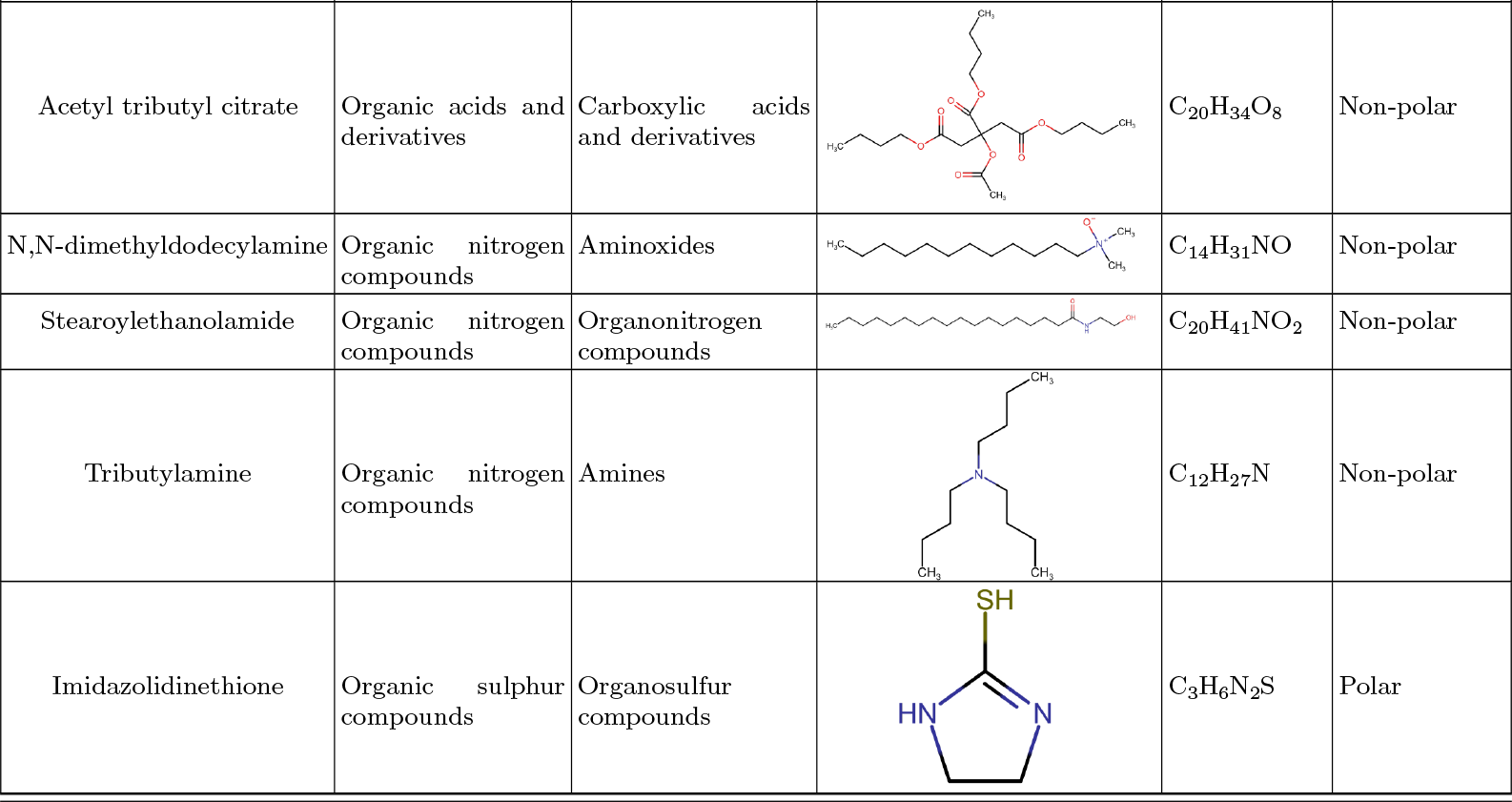
Classification of identified metabolites. Classifications obtained from Global Natural Products Social Molecular Networking [54] (GNPS), Metabolomics Workbench [61]. Polar identifications were previously described in [19].

