## Supplementary Materials (Tables S1, S2) for "Early viral infection of cyanobacteria drives bacterial chemotaxis in the oceans"

### Early stage viral infection of cyanobacteria drives marine bacterial chemotaxis

This Supplementary Information file includes:

Supplementary Table S1: **Mass spectrometer parameters**

Supplementary Table S2: **RP chromatography**

Supplementary File: **SupplementaryData_S3.xlsx**

Excel spreadsheet containing Supplementary Table S3: Values and significance from metabolomic analysis/compound identification

Supplementary File: **SupplementaryData_MZmine2-params-PhageInfection.zip**

Compressed folder containing parameters used (in batch format) with MZmine 2.39 for feature detection

Table S1: Mass spectrometer parameters

| **Mass Spectrometer Parameters** | |
|---|---|
| Instrument | Thermo Q Exactive |
| Data acquisition mode | Full MS -> ddMS2 |
| Ion polarity mode | Injected 2x: first in positive mode, then in negative mode |
| **Source Settings** | |
| Sheath gas flow rate | 55 |
| Aux gas flow rate | 20 |
| Sweep gas flow rate | 2 |
| Spray voltage ( \|kV\| ) | 3 |
| Capillary temp (degrees C) | 400 |
| S-lens RF level | 50 |
| **Full MS Settings** | |
| Resolution | 70000 |
| AGC target | 3.00E+06 |
| Max. IT | 100ms |
| Scan range | 70-1050 m/z |
| Spectrum data type | Centroid |
| **dd-MS2 Settings** | |
| Resolution | 17500 |
| AGC target | 1.00E+05 |
| Max. IT | 50 ms |
| Loop count | 2 |
| TopN | 2 |
| Isolation window (m/z) | 1 |
| (N)CE/stepped nce | 10,20,40 |
| Spectrum data type | Centroid |
| **dd Settings** | |
| Min. AGC target | 1.00E+03 |
| Charge exclusion | >3 |
| Exclude isotope | on |
| Dynamic exclusion | 7 seconds |
| **LC parameters** | |
| LC stack | Agilent 1290 UHPLC |
| Autosampler temperature | 4 degrees C |
| Injection volume | 4 uL |
| Injection draw and eject speeds | 20 uL/min @ 2mm draw position |
| Additional injection settings | 2 second equilibration time and 5x flush out factor |
| Needle wash | 3 seconds, mode = flush port using 50:50 v/v methanol:water |
| Control notes | Internal (in all samples) and external (injected every 10 samples) standards were used for quality control purposes, methanol blanks injected between samples |

Table S2: RP chromatography

| RP chromatography | |
|---|---|
| Column | Agilent 959757-902 ZORBAX RRHD Eclipse Plus C18, 95Å, 2.1 x 50 mm, 1.8 µm, 1200 bar pressure limit |
| Column model | 959757-902 |
| Column serial number | USDAY46920 |
| Composition of mobile phase solvent A | 0.1% v/v formic acid in water |
| Composition of mobile phase solvent B | 0.1% v/v formic acid in acetonitrile |
| Column compartment temperature | held at 50 degrees C |

| Timetable | | | |
|---|---|---|---|
| **Time (min)** | **Flow (ml/min)** | **%A** | **%B** |
| 0 | 0.4 | 100 | 0 |
| 1 | 0.4 | 100 | 0 |
| 8 | 0.4 | 0 | 100 |
| 9.5 | 0.4 | 0 | 100 |
| 10.5 | 0.4 | 100 | 0 |
| 11.5 | 0.4 | 100 | 0 |

| Internal standards | | | |
|---|---|---|---|
| **Component name** | **Vendor** | **Product SKU** | **Concentration** |
| 13C,15N Cell Free Amino Acid Mixture | Sigma-Aldrich | 767964-1EA | diluted 1:2000 |
| 13C-a,a,-trehalose | Omicron | TRE-002 | 10 ug/mL |
| 13C-mannitol | Omicron | ALD-030 | 10 ug/mL |
| 2-amino-3-bromo-5-methylbenzoic acid | Sigma-Aldrich | R435902-250MG | 1 ug/mL |
| 13C-15N-cytosine | Sigma-Aldrich | 492108-10MG | 5 ug/mL |
| 13C-15N-uracil | CIL | CNLM-3917-PK | 2 ug/mL |
| 15N-inosine | CIL | NLM-4264-PK | 5.5 ug/mL |
| 15N-adenine | CIL | NLM-6924-PK | 4 ug/mL |
| 15N-hypoxanthine | CIL | NLM-8500-PK | 3 ug/mL |

| Identification details |
|---|
| Untargeted: Most stereoisomers and some positional isomers are not resolvable on the methods used. Our knowledge of isomer overlaps is limited by the availability of reference standards in our RT and MSMS library, thus other isomers that are not listed may also be possible; if untargeted features matched to online MSMS databases for identification, these will not be compared with known RT and may reflect an incorrect isomer match. |
